# Ethanol exposure model in zebrafish causes phenotypic, behavioral and gene expression changes that mimic Fetal Alcohol Spectrum Disorders in human birth cohorts

**DOI:** 10.1101/2024.08.05.606708

**Authors:** Elanur Yilmaz, Nastasia Nelson, Giuseppe Tosto, R. Colin Carter, Caghan Kizil

## Abstract

Fetal Alcohol Spectrum Disorders (FASD) represent a significant global health challenge, characterized by physical and neurodevelopmental abnormalities in offspring resulting from prenatal alcohol exposure. This study aims to utilize the zebrafish to examine the phenotypic, behavioral, and molecular changes associated with embryonic ethanol exposure, providing a model for human FASD conditions. Our study exposed zebrafish embryos to 0.5% ethanol during a critical developmental window (2-24 hours post-fertilization) and documented significant craniofacial and cardiac deformities, which recapitulate what has been observed in human FASD in humans. Notably, exposed zebrafish exhibited reduced skull and eye sizes, thickened jaw size, and enlarged heart chambers. We found reduced burst swim distance following a touch stimulus, a novel behavioral assessment of potential deficits in sensory processing such as processing speed and/or stress/startle response, both of which are affected in human FASD. Whole-organism gene expression was found to be altered by ethanol for orthologs of four of five inflammation-related genes for which placental expression was previously found to be altered in response to alcohol in human placentas (*SERPINE1, CRHB, BCL2L1, PSMB4, PTGS2A*). We conclude that the zebrafish model effectively mimics several FASD phenotypes observed in humans, confirming gene expression changes we have previously documented in a human observational study and providing a valuable platform for exploring the underlying mechanisms of alcohol-induced embryonic alterations and for developing diagnostic markers and therapeutic targets for early intervention.

## INTRODUCTION

Fetal alcohol spectrum disorders (FASD) are one of the foremost preventable causes of neurodevelopmental disabilities worldwide. Prevalence estimates range from 1-5% in the US and Western Europe (May et al., 2009) to 13.6-30.6% of school-aged children in endemic communities in South Africa (May et al., 2021). FASD manifest a broad range of cognitive and behavioral deficits and growth restriction, and teratogenic effects of alcohol have been reported in every organ system (Carter et al., 2016a; Carter et al., 2012; Carter et al., 2016b; Hoyme et al., 2005; Jacobson, 1998). Despite behavioral intervention programs and extensive prevention efforts (ACOG, 2011), ∼18% of US women continue to drink during pregnancy, with 6.6% reporting binge-drinking episodes (SAMHSA, 2010). Postnatal treatment programs to improve cognitive abilities in children with FASD are limited by difficulties in identifying which children to target. While fetal alcohol syndrome (FAS) is characterized by specific facial dysmorphology and growth restriction (Hoyme et al., 2016; Hoyme *et al*., 2005), children with the most prevalent form of FASD, alcohol-related neurodevelopmental disorder (ARND), are difficult to identify because they lack the characteristic facial dysmorphology and are thus often identified later in life when early intervention approaches are less effective. Yet these children still have cognitive impairments as severe as those with FAS. Thus, improved diagnostic tools are needed, such as more fully characterized behavioral phenotypes and molecular biomarkers of effect, e.g., gene expression profiles detectable at early life stages that identify children with teratogenic effects of alcohol exposure. In prospective longitudinal birth cohorts in Cape Town, South Africa, we helped to characterize the evolution of characteristic FASD dysmorphology across childhood and adolescence (Jacobson et al., 2021), fetal alcohol-related neurobehavioral deficits (Carter et al., 2022; du Plessis et al., 2015; Fan et al., 2016; Jacobson et al., 2011), including slower processing speed, and alcohol exposure-related alterations in -placental expression of genes related to growth, inflammation, and neurobehavior (Carter et al., 2018; Deyssenroth et al., 2024; Masehi-Lano et al., 2023; Williams et al., 2023).

As these and other FASD studies in humans are observational, animal models are needed to validate and functionally characterize such findings. Zebrafish serve as a unique vertebrate experimental model system, as they have unusually high morphological and genomic conservation with humans and experimental mutability of embryogenesis and development that is mostly found in invertebrate animal models, such as *C. elegans* or *D. melanogaster* (Adhish and Manjubala, 2023; Burgess and Burton, 2023). Furthermore, the zebrafish genome and developmental transcriptome are well-characterized, with known timing of gene expression patterns/profile and a well-described transition from expression of maternal genes in the egg to embryonic genes in the maternal-zygotic transition in early development (Burgess and Burton, 2023; White et al., 2017). Thus, the timing of teratogenic insults and their effects can be identified in experimental models. Accordingly, FASD zebrafish models have led to foundational contributions to understanding of the pathophysiological underpinnings of the teratogenic effects of alcohol. Zebrafish models have robustly replicated structural effects of ethanol exposure on craniofacial, ocular, brain, and cardiac embryologic development and have helped to elucidate genetic and molecular underpinnings of these effects (Adhish and Manjubala, 2023; Fernandes and Lovely, 2021). Zebrafish models have also helped to characterize alcohol-related behavioral deficits in social behavior, learning and memory, and anxiety phenotypic outcomes (Adhish and Manjubala, 2023; Fernandes and Lovely, 2021).

In our FASD zebrafish model, we sought to replicate findings from our longitudinal human birth cohorts, focusing on alcohol-related effects on craniofacial development, processing speed, and gene expression. Such replication would validate the findings from our observational studies and establish readouts in FASD zebrafish model that can be used in future experiments to identify genetic and molecular mechanisms underlying these teratogenic effects of alcohol and potentially test therapeutic/preventive treatments tailored to such mechanisms.

## RESULTS

We implemented a zebrafish model to investigate the effects of embryonic ethanol exposure, aiming to replicate and expand upon findings from human longitudinal birth cohorts that documented craniofacial development alterations, processing speed deficits, and inflammation-related gene expression changes due to prenatal alcohol exposure. We subjected zebrafish embryos to 0.5% ethanol exposure from 2 to 24 hours post-fertilization and conducted touch evoked response assay at three days post-fertilization (**Figure 1A**). Morphological measurements were done on day 4.5 to evaluate the overall body plan properly.

**Figure 1:**
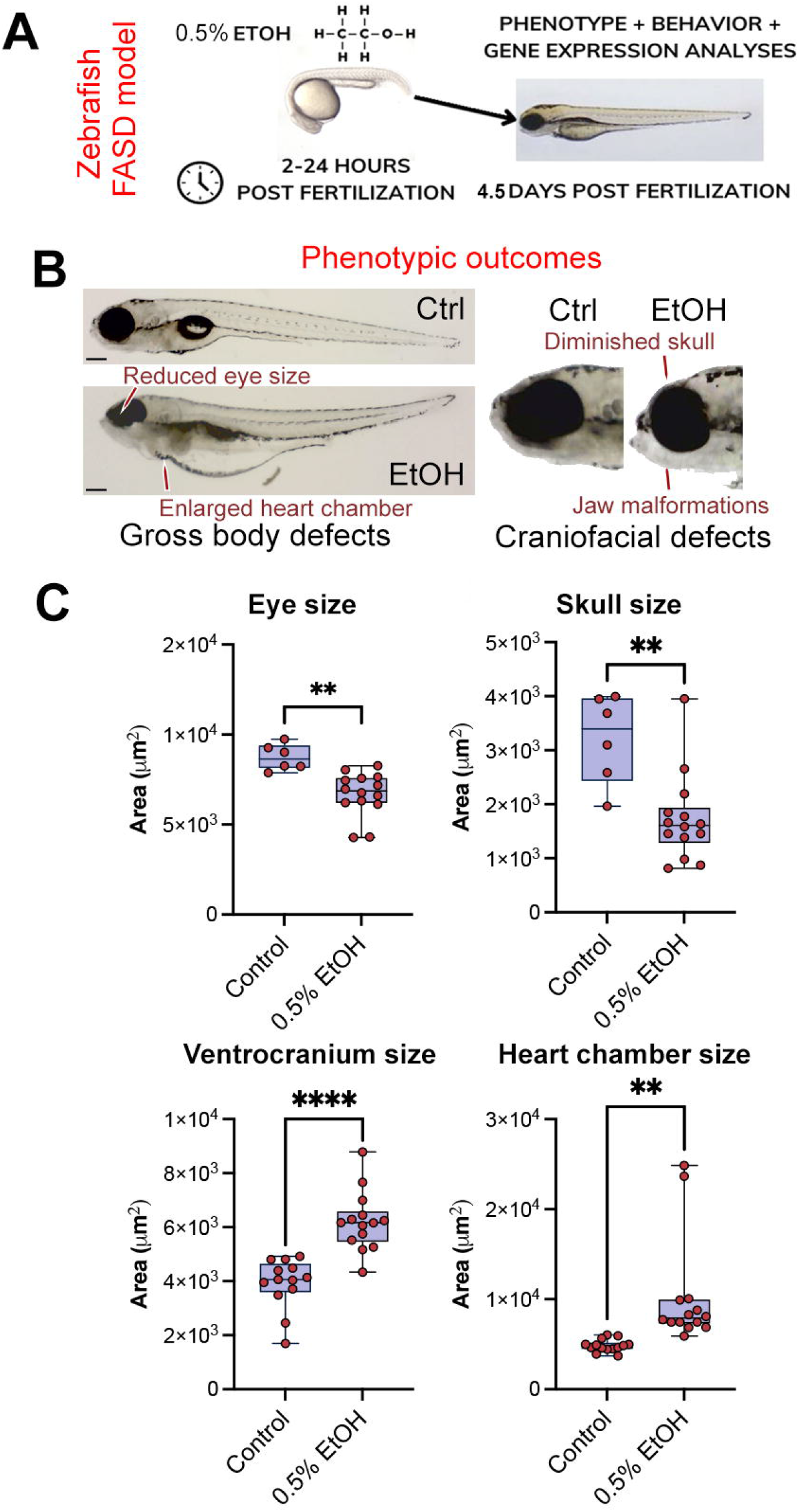
Ethanol exposure causes phenotypic changes. (A) Schematic view of the FASD model. (B) Exemplary phenotypic changes in zebrafish upon ethanol exposure. (C) Quantification of the sizes of eye, skull, ventrocranium, and heart chamber in control and 0.5% ethanol-treated groups. Scale bars equal 100 microns. *p*<0.001: **, *p*<0.0001: ****.

### Phenotypic changes with alcohol exposure

After ethanol exposure in zebrafish, we found significant craniofacial and cardiac abnormalities in the ethanol-exposed group (FASD) compared to controls (**Figure 1B, 1C**). In the sagittal plane, skull size was reduced in ethanol treated animals when compared to control animals [1.7 vs. 3.2 mm^2^, *t*(18)=3.743, *p*=0.0015]. Eye size was similarly reduced (6.7 vs. 8.7 mm^2^, *t*(18)=3.807, *p*=0.0013). To examine potential alterations in cranio-facial cartilage development, we performed Alcian blue staining in control and 0.5% EtOH-treated animals (**Figure 2A**) and observed significant alterations in jaw formation (**Figure 2B**). Ethanol treatment led to reduced distance between Meckel’s cartilage (MK) and ceratohyal cartilage (CH) [101.81 μm vs. 85.29 μm, *t*(53)=3.57, *p*=7.75E-04], reduced length of MK [1,376.95 μm vs. 973.61 μm, *t*(48)=6.229, *p*=1.2E-07], and reduced dorsoventral cartilage thickness [172.71 μm vs. 106.50 μm, *t*(24)=4.497, *p*=1.49E-04]. The angle between CH formations was not altered [85.65 vs. 83.61 degrees, *t*(53)=0.919, *p*=0.3622]. This constellation of findings indicates disrupted chondrogenesis and growth patterns of the craniofacial skeleton consistent with facial hypoplasia. Ethanol treatment was also associated with an enlarged heart chamber (**Figure 1C;** 10.2 vs. 4.8 mm^2^ in sagittal section, *t*(26)=3.340, *p*=0.0025).

**Figure 2:**
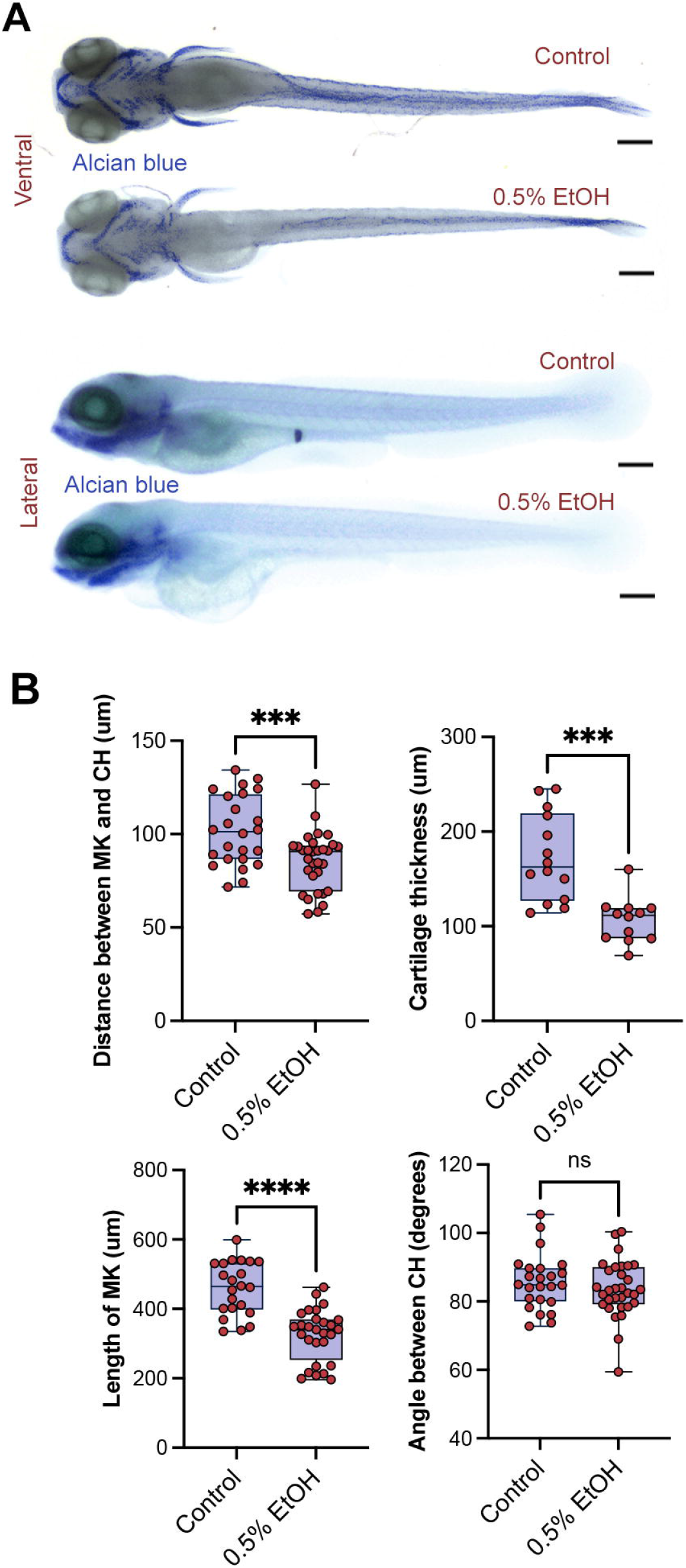
Ethanol exposure causes facial hypoplasia. (A) Alcian blue staining of jaw cartilage. (B) Unpaired parametric *t*-test comparisons of jaw measurements. Scale bars equal 100 microns. *p*<0.0001: ***, *p*<0.0001: ****.

### Alterations in behavioral and locomotor functions after alcohol exposure

To determine whether alcohol exposure alters response to a stressful stimulus and the locomotor activity, we performed needle tail touch stimulus at 3 days after fertilization, in which a needle touches the tail of the animal causing a normative flight response (**Figure 3A, B**). The distance swum and the speed at which swimming occurs are both recorded. The burst swim response following a touch stimulus - a primitive reflex that is largely conserved across many species, including zebrafish and human - is a critical measure for the rapid escape behavior that relies on the proper processing of the environmental cue in the spinal cord and brain (Koutsikou et al., 2018). We found that ethanol-treated zebrafish displayed a decreased burst swim distance in response to a needle touch stimulus when compared to control animals (9.4 vs. 14.3 mm, *t*(10)=2.370, *p*=0.0393), indicating a diminished response to this stressful stimulus (**Figure 3C**). By contrast, swimming speed was unchanged (125.0 vs. 152.1 mm/second, *t*(10)=-0.73, *p*=0.483) (**Figure 3C**).

**Figure 3:**
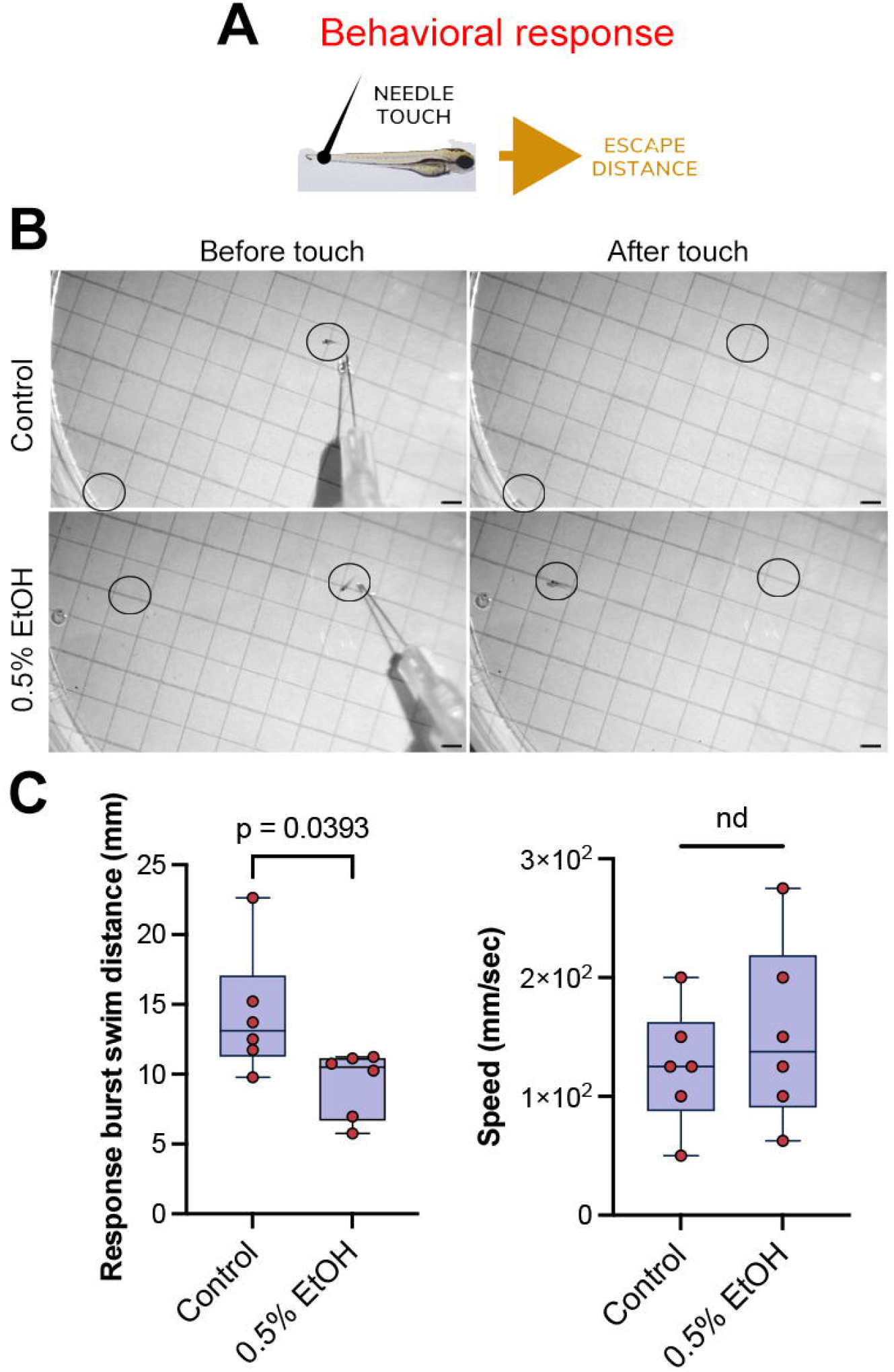
Ethanol exposure alters stimulus processing. (A) Schematics for needle touch stimulus assay. (B) Representative images of recordings. Circles are start and end points after one round of tail touch. Scale bars equal 1 mm. (C) Quantification of burst swim distance and swimming speed of escape after stimulus (unpaired parametric *t*-test comparisons). Scale bars equal 1 mm.

### Gene expression changes in alcohol-exposed zebrafish

We also examined whole-organism gene expression across a range of ethanol exposures for *serpine1, crhb, bcl2l1, psmb4*, and *ptgs2a*, zebrafish orthologs of inflammation-related genes found to be differentially expressed upon alcohol exposure in placenta samples in our human birth cohort (Deyssenroth *et al*., 2024; Williams *et al*., 2023). Recapitulating our human placental findings, ethanol treatment was related to altered expression of four genes (**Figure 4**). For *serpine1*, a positive linear dose-response pattern was seen from ethanol concentrations of 0.5%, the concentration that most closely approximates human blood alcohol levels associated with FASD, and higher. For *crhb*, a negative linear dose-response pattern was seen from ethanol concentrations of 1% and higher. For *bcl2l1* and *psmb4*, gene expression was lower among animals with the highest exposure (2%) compared with unexposed control animals. Only *ptgs2a* showed no differential expression in response to staggered alcohol doses.

**Figure 4:**
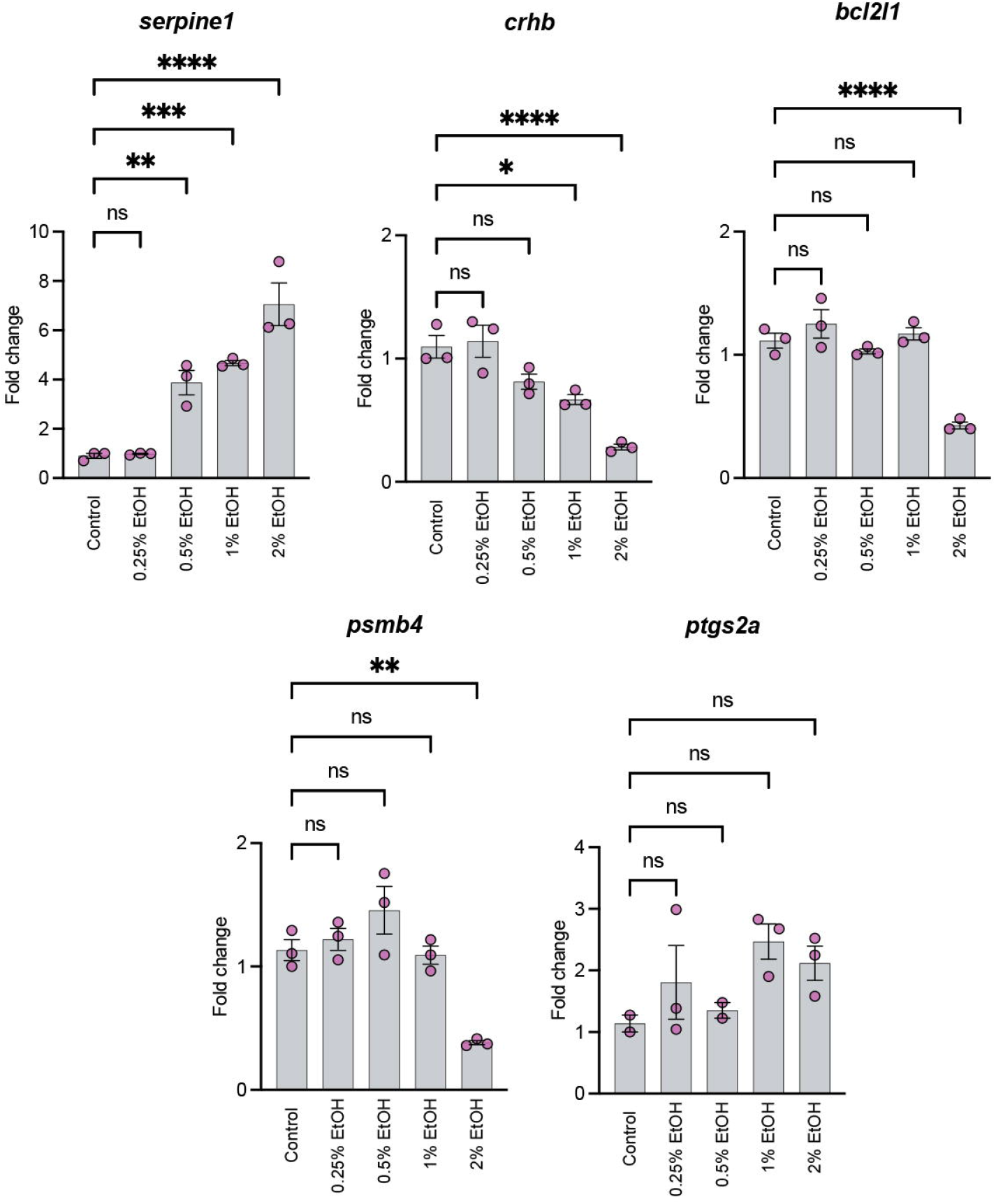
Gene expression comparisons using quantitative real-time PCR for 5 genes differentially expressed based on prenatal alcohol exposure in human placentas. Quantification graphs after different doses of alcohol exposure in zebrafish embryos. ANOVA and Dunnett’s post-hoc test. *p*<0.032: *, *p*<0.0021: **, *p*<0.0002: ***, *p*<0.0001: ****.

## DISCUSSION

In our FASD zebrafish model, we demonstrated the effects of embryologic ethanol exposure on craniofacial development, behavioral response to a needle touch stimulus, and expression of genes with roles in neuroinflammation, a key mechanism underlying ethanol-induced brain damage. These findings are consistent with prior findings in human studies (Carter *et al*., 2018; Carter *et al*., 2022; Deyssenroth *et al*., 2024; du Plessis *et al*., 2015; Fan *et al*., 2016; Jacobson *et al*., 2021; Jacobson *et al*., 2011; Masehi-Lano *et al*., 2023; Williams *et al*., 2023) and establish an experimental model for future examination of genetic and molecular pathways underlying the effects of alcohol on these outcomes.

The ethanol-induced embryologic structural changes we demonstrated validate our FASD zebrafish model and experimental paradigm, as such anomalies parallel previous zebrafish models and well-documented phenotypes associated with FASD in humans, including reduced head circumference and characteristic ocular dysmorphologies (Cadena et al., 2020; Fernandes et al., 2014; Sarmah et al., 2016; Sidik et al., 2021). As expected from prior FAD zebrafish models (Alsakran and Kudoh, 2021; Fernandes and Lovely, 2021), embryologic ethanol exposure from 2-24 hours post-fertilization led to reductions in skull and eye size, cranial hypoplasia, and enlargement of the heart chamber. Embryonic reductions in skull size are the result of smaller brain size, and the human analogue, reduced head circumference, is one of the key diagnostic criteria for FAS and partial FAS (Hoyme *et al*., 2016; Hoyme *et al*., 2005; Jacobson *et al*., 2021). Previous zebrafish experiments have demonstrated ocular defects in ethanol-exposed zebrafish, including reduced eye size, as we have shown here (Dlugos and Rabin, 2007). In humans, fetal alcohol effects on the visual system are well-documented (Carter et al., 2005; Stromland, 1987), and ocular dysmorphic features are characteristic of FASD dysmorphology, including inner- and outer-canthal distances, interpupillary distance, and, most notably, palpebral fissure length, i.e., the measure of the length of the eye opening (Gomez et al., 2020; Jacobson *et al*., 2021). Indeed, reduced palpebral fissure length is one of the three hallmark dysmorphic features used in FASD diagnostic criteria (Hoyme *et al*., 2016; Hoyme *et al*., 2005). Embryologically, reductions in palpebral fissure length are thought to be driven by reduced ocular size and alterations in forebrain development, consistent with our findings. Our finding of ethanol-associated facial hypoplasia has been previously shown in zebrafish models (Sidik *et al*., 2021), and, in the midface, is well-characterized in humans (Muggli et al., 2017; Suttie et al., 2013; Suttie et al., 2017). Lastly, although the cardiotoxic effects of prenatal alcohol exposure in humans are more variable and less well-characterized than craniofacial dysmorphology, our finding of ethanol-induced heart chamber enlargement is consistent with prior zebrafish models (Dlugos and Rabin, 2007).

These morphological changes observed in our zebrafish model reflect disruptions in developmental processes that are critical for normal organogenesis. Ethanol exposure might be interfering with several key pathways. Alcohol inhibits the proliferation of neural crest cells and disrupt their migration patterns (Smith et al., 2014). Neural crest cells are essential for the development of craniofacial structures and the heart (Achilleos and Trainor, 2012). Impaired migration and differentiation of these cells due to ethanol exposure could lead to the craniofacial abnormalities and cardiac defects we observed. Additionally, ethanol might affect the expression and function of genes critical for development. This includes genes involved in the growth and differentiation of cells in the craniofacial region and heart. It is known that ethanol metabolism produces reactive oxygen species (ROS), leading to oxidative stress which can damage cells and tissues, impairing normal development (Qin and Crews, 2012). This oxidative stress is particularly detrimental during the rapid growth and differentiation stages of early embryogenesis as it can induce apoptosis in developing tissues (Redza-Dutordoir and Averill-Bates, 2016). The correlation of the phenotypes in zebrafish to humans not only underscores the utility of the zebrafish model in mimicking human FASD phenotypes but also provides a controlled environment to further explore the specific mechanisms by which alcohol alters embryonic development.

We also demonstrated ethanol-induced reductions in burst swim distance seen after a needle-touch stimulus to the animal’s tail. This finding is particularly significant as it represents a novel behavioral assessment in the FASD zebrafish model. To our knowledge, no prior study has examined this outcome in an FASD zebrafish model. The observed deficit in burst swim distance may reflect diminished sensory processing, slower processing speed, diminished startle response, and/or gross locomotor deficits. Interestingly, despite these reductions in burst swim distance, our study did not find significant changes in gross locomotor function as indicated by swimming speed (**Figure 2C**). This suggests that the primary effect of ethanol exposure may be on the sensory and neural processing pathways rather than on the basic motor functions.

The ability to respond to a needle touch with a rapid swimming motion also involves sensory processing (Abraira and Ginty, 2013), which is coordinated by multiple neural circuits, including sensory detection, signal processing in the nervous system, and motor execution. In a normative response, the sensory systems must detect the stimulus, the nervous system must process this information, and the motor systems must execute the escape response. Impairments in any part of this sensory integration pathway can result in a diminished response. The diminished burst swim response indicates that prenatal ethanol exposure could impair these critical neural pathways. This is consistent with human studies showing that individuals with FASD often have slower processing speed and impaired startle responses, which can affect their daily functioning and response to environmental stimuli. We and others have demonstrated prenatal alcohol exposure-associated processing speed deficits in human studies (Burden et al., 2009; Carter *et al*., 2022; Fan et al., 2017; Jacobson, 1998; Jacobson et al., 1994; Willford et al., 2010), which may have pervasive functional relevance in activities of daily living and function in employment for affected individuals.The burst swim is part of the startle response, a fundamental survival mechanism. This response can also be influenced by the hypothalamic-pituitary-adrenal (HPA) axis. Animal models and human studies have demonstrated fetal alcohol-induced alterations in the hypothalamic-pituitary-adrenal (HPA) axis and stress-responses, with blunted behavioral and biochemical responses seen early in life and overactive responses and related mental health issues later in life (Fifer et al., 2009; Hellemans et al., 2010; Oberlander et al., 2010; Ruffaner-Hanson et al., 2022). Our findings underscore the utility value of using zebrafish as a model to study the specific neural and molecular mechanisms underlying these behavioral deficits. Future studies should focus on further characterizing the ethanol-induced changes in sensory processing and startle response pathways. Investigating the neural circuits and gene expression profiles involved in these behaviors could provide deeper insights into how prenatal alcohol exposure disrupts brain development and function. Moreover, understanding these mechanisms may open avenues for developing targeted therapeutic interventions to mitigate the effects of FASD on sensory and neural processing.

A growing body of literature has implicated neuroinflammation in fetal alcohol-related brain damage (Kane and Drew, 2021). For example, in a rat model, prenatal alcohol exposure led to higher levels of cytokines and chemokines in the fetal brain and the placenta, higher cytokine/chemokine response to lipopolysaccharide stimulation in the adult offspring brain, and related memory impairments (Terasaki and Schwarz, 2017). Our findings of ethanol-induced alterations in *serpine1, crhb, bcl2l1*, and *psmb4* gene expression are consistent with our recent reports of alcohol-related alterations of these and other inflammation-related genes in alcohol exposed placenta samples from a human longitudinal birth cohort (Deyssenroth *et al*., 2024; Williams *et al*., 2023). These findings validate our prior findings in humans and identify pathways that can be explored for future studies of targeted therapeutics. In the current study, a positive dose-response relationship was seen between ethanol exposure and zebrafish *serpine1* expression. These findings are consistent with our finding of alcohol exposure-related increases in placental expression of the human analogue, *SERPINE1*, which encodes a member of the serine proteinase inhibitor (serpin) superfamily that plays roles in antiviral immunity and contributes to regulation of neuroinflammation (Pelisch et al., 2015; Pu et al., 2022; Wang et al., 2021). In our zebrafish model, ethanol was associated with decreased *crhb* expression. *crhb* in zebrafish and its analogue *CRH* in humans are key regulators of the hypothalamic-pituitary-adrenal (HPA) axis; disruptions in the HPA axis have been repeatedly demonstrated to play roles in FASD inflammation-related pathophysiology in both animals and humans (Ruffaner-Hanson *et al*., 2022). Interestingly, while *crhb* expression was decreased in response to ethanol in our zebrafish model, CRH was increased in our human placenta study. This difference may be due to differences between species (e.g., zebrafish do not have placentas). Notably, prior FASD zebrafish models have shown both up- and down-regulation of *crhb* in response to developmental ethanol exposure, depending on the developmental timing of the exposure and the age at which *crhb* expression was measured (Du et al., 2020). The ethanol-induced reductions in *bcl2l1* seen in our zebrafish model match our finding in human placentas and prior work demonstrating decreased hepatic expression of *BCL2L1* in ethanol-exposed rats (French et al., 2005) and decreased expression in an ethanol-treated mouse hippocampal cell line (HT22) (Song et al., 2014). The human analogue *BCL2L1* is a regulator of apoptosis via reactive oxygen species production and cytochrome C release in mitochondria. Its isoform Bcl-X(L) also regulates presynaptic activity, neurotransmitter release/recovery, as well as synaptic vesicle formation and endocytic vesicle retrieval in the hippocampus. In an immune function role, it may impair NLRP1-inflammasome activation (Bruey et al., 2007), which is notable since bcl2l1 was down-regulated in ethanol exposed animals in the current study. *psmb4* expression was decreased in response to ethanol exposure, consistent with our finding in human placentas of alcohol-related reductions in expression of *PSMB4*, which encodes a proteosome subunit important for the immunoproteosome that facilitates non-lysosomal, ubiquitin-independent peptide cleavage necessary for MHC class I-presented antigenic peptides (Verhoeven et al., 2022). Although the mean fold change expression levels for *ptgs2* were higher in the 1% and 2% ethanol treatment groups than for the lower concentration groups, these differences were not statistically significant. *PTGS2* encodes the inducible form of prostaglandin-endoperoxide synthase, a.k.a., cyclooxygenase, a key enzyme in prostaglandin synthesis that plays roles in the creation of resolvins (resolution phase interaction products) during inflammation (Serhan et al., 2002) and in neuronal secretion of specialized preresolving mediators (SPMs) that regulate phagocytic microglia. The mouse analogue, *Ptgs2*, has been shown to play key roles in neuroinflammation regulation in cerebral ischemia (Jiang et al., 2023) and traumatic brain injury (Graber et al., 2015).

While our model offers substantial insights, it has limitations. The controlled environment of our study may not fully encapsulate the complexity of alcohol metabolism in a pregnant human, where varying factors such as genetics, overall health, nutritional status, and additional substance use can significantly affect the teratogenic outcomes. Our study did not examine potential cumulative effects of repeated or prolonged exposure to ethanol, as might be more typical in human scenarios of alcohol use during pregnancy. Future studies could look to model different patterns of exposure to better replicate the range of exposure scenarios seen in human populations. Early developmental models do not answer the long-term effects of early alcohol exposure. Longitudinal studies within our zebrafish model could provide further insights into how early alcohol exposure influences adult phenotypes, particularly in neurobehavioral contexts.

Overall, our findings demonstrate that the zebrafish FASD model effectively replicates and extends critical aspects of human FASD, including morphological changes, behavioral deficits, and molecular alterations. This model not only helps in understanding the pathophysiological basis of FASD but also provides a viable platform for testing potential interventions and studying the underlying molecular mechanisms in a controlled environment. We replicated fetal alcohol-related effects on craniofacial and heart development, behavior, and gene expression seen in prospective longitudinal birth cohorts. Furthermore, we established a new FASD zebrafish model, which includes a novel alcohol-related needle-touch behavioral deficit, consistent with processing speed and/or stress-response changes in human studies. Our FASD zebrafish model may be used in future work to examine how specific genetic factors, identified in human studies, influence fetal development in the setting of ethanol exposure and could uncover critical periods of vulnerability to ethanol, thus enhancing our understanding of FASD susceptibility. Functional characterization of genetic variants associated with FASD could elucidate the molecular mechanisms driving this disorder. This could involve assessing how these variants impact gene expression, protein function, and cellular pathways. For example, employing CRISPR/Cas9 technology to edit genes implicated in FASD within this model could validate the roles of specific genes and uncover potential therapeutic targets.

## MATERIALS AND METHODS

### Zebrafish husbandry

Adult animals were maintained under temperature and humidity-controlled environment of Columbia University Medical Center on a 14/10-h light/dark cycle, with ad libitum access to food and water in compliance with the protocols approved by the Institutional Animal Care and Use Committee of Columbia University (IACUC). All zebrafish experiments were performed in larval stages (up to day 5) in accordance with the applicable regulations by the IACUC. Adult male/female zebrafish with a 3:3 ratio was isolated in slope breeding tank with a divider. The next morning, dividers were removed, and the eggs were collected 30 minutes after crossing. The eggs were collected and incubated in 28.5°C incubator. 2 hours later fertilized eggs with normal developing stage were randomly selected and distributed to the experimental groups, while unfertilized eggs with abnormal development and/or malformations were discarded. The survival rate of animals in the control group at day 1 was ≥ 95% and the hatching rate was 100% at day 3.

### Ethanol preparation and treatment

Absolute ethanol (Lot: 222257, CAS # 64-17-5, Fisher Bioreagents) was used to prepare 0.25%, 0.5%, 1%, 2% ethanol concentrations. EtOH solution prepared freshly before the experiment and added to the embryo medium. We followed the methodology of FASD models as described before (Cadena *et al*., 2020; Sarmah *et al*., 2016). In short, the healthy embryos were raised in embryo medium with methylene blue until 2 hour-post-fertilization (hpf). Developmental stages were defined as described (Kimmel et al., 1995; Kimmel et al., 1988). At 2 hpf, equal number of embryos were transferred to the embryo medium without methylene blue (untreated, control group) and medium with related EtOH concentrations (0.25% (∼50mM), 0.5% (∼100mM), 1% (∼200mM), 2% (∼400mM) in Petri dishes by wrapping with Parafilm^®^. Following the 22 hours of exposure (at 24 hpf), all solutions were replaced with fresh embryo mediums with methylene blue. All experimental groups were raised in 28.5°C incubator until the experiment.

### Behavioral measurements

Embryos that are dead or have severe morphological defects (≤ 9% for all groups) were excluded from the behavioral test. In the end we analyzed the touch evoked response in 6 embryos in each group. The larvae were placed in a large petri dish under grid paper (0.50 cm per gridbox). A dissecting scope attached digital camera (AmScope, SM-4T) was elevated 12 inches to enable a wider view of the larvae in the dish. One larva at a time was transferred into the dish using a Pasteur pipette. After acclimation, the larvae were prodded with a fine point syringe needle on the tail until an attempted to escape was prompted, and swimming response were recorded.

### Video analysis

The larvae videos were analyzed utilizing iMovie. The number of needle touches before an escape attempt was recorded. The Start Time was considered the last needle touch before larva locomotion (or larva’s escape attempt). The End Time was the first stop made by the larva after the escape attempt. The distance travelled was calculated by multiplying 0.50 cm by the number of grid boxes traveled. The rate of travel was evaluated by subtracting the start time from the end time and dividing it by the distance travelled. Additionally, we utilized iMovie’s freeze frame and slow-motion features to accurately capture the above perimeters and straight path criteria. The straight path variable was defined as the larva changing direction after proceeding on a specific path.

### Alcian Blue staining

Alcian blue staining was performed [as described in (Neuhauss et al., 1996)] to 4 dpf zebrafish larvae of control and 0.5% EtOH treated group to detect craniofacial cartilage morphology. Briefly, zebrafish larvae were fixed in 4% PFA in PBS overnight in cold room. The rest of the steps were performed at room temperature, and on the shaker. After fixation, larvae were washed three times with PBST, each washing was for 5 min. Larvae then were dehydrated by using 50% ethanol for 15 min and stained overnight with Alcian Blue (Sigma, Cat no: B8438). The next day, larvae were washed with PBST for 5 min and bleached (30% H_2_O_2_ (Sigma, Cat no: H1009) and 2% KOH (Sigma, Cat no: 417661)) for 1 hour. After removing the bleach solution, larvae were washed for 5 min with PBST and cleared with 25% Glycerol (Sigma, Cat no: G5516) contains 0.25% KOH and stored in 50% Glycerol with 0.25% KOH until imaging with Zeiss Discovery. V12 with AxioCam MRc and AxioVision LE64 software.

### Morphological analysis

At day 4.5, control and 0.5% EtOH-treated experimental groups were analyzed by their craniofacial (eye distance, eye and skull size, jaw position), cardiac (heart chamber size), and overall morphological phenotypes (body length, spine deformation, tail deformation, yolk sac edema, swim bladder inflation). For craniofacial cartilage analysis, distance between rostromedial regions of Meckel cartilage (MK) and ceratohyal cartilage (CH) (on ventral views), length of the MK (on ventral images), angle between lateral CH cartilages (on ventral images), and the cartilage thickness below the lens (lateral images) were measured using ImageJ software 2.1.0/v1.53.c.

### Quantitative real-time PCR

We examined zebrafish orthologs for 5 candidate inflammation-related genes found to be differentially expressed based on alcohol exposure (control (n=11), 0.25% (n=11), 0.5% (n=12), 1% (n=12) and 2% (n=12)) in human placentas in our prospective birth cohort (Deyssenroth *et al*., 2024; Williams *et al*., 2023). RNA isolation of pooled embryos of each group was performed by using TRIzol^®^ Reagent (Invitrogen, 15596026) and Direct-zol RNA Microprep Kit (Zymo Research, R2060) according to the manufacturers’s instructions. cDNA was synthesized from the purified RNA using the SuperScript™ III First-Strand Synthesis System (18080051; Invitrogen; Thermo Fisher Scientific, Inc.) with oligo (dT) by following the manufacturer’s protocol. The qRT-PCR reaction was performed using a final volume of 10 μl includes 5 μl of PowerUp™ SYBR™ Green Master Mix 2x (Applied Biosystems, A25742), 1 μM of each primer pair (*serpine1* forward: GCCGTTAACTGTGTGGAAGA, *serpine1* reverse: ACGGTTAGTCTTCCTGAATGC; *crhb* forward: CGTAACTGCCATCCAAGCG, *crhb* reverse: TCCCCGAGACATCCCAGTAT; *bcl2l1* forward: CTCCTTCTCCACACACTCGA, *bcl2l1* reverse: GCGTGATGGATGAGGTGTTT; *psmb4* forward: GGATGGTGCTGTTGTTGACT, *psmb4* reverse: GTTCAAGGGTGGTGTGATCA; *ptgs2a* forward: GATGAGTAGGGCTTCATGTTGA, *ptgs2a* reverse: CACCTGCAGTTCAAGGAGTC; *β-actin* forward: ATGCAGAAGGAGATCACATCCC, *β-actin* reverse: GCTTGCTAATCCACATCTGCTG), 1 μl of the cDNA (10 ng in final concentration) and 2 μl of UltraPure™ DNase/RNase-Free Distilled Water dH2O (Invitrogen™ Cat.#.: 10977015). The cycle threshold (Ct) value for the target genes was calculated using QuantStudio™ 3 real-time PCR system. Amplification conditions for qRT-PCR were as follows: 50 °C for 2 minutes, 95°C for 10 minutes, followed by 40 cycles at 95 °C (15 seconds) and 60 °C (1 minute). Melting curve was analyzed for each sample by following 95 °C for 15s, 60°C for 1 minute and 95 °C for 1 second after amplification reactions. Relative expression levels were calculated using ΔΔCt method.

## Statistical analyses

Pairwise comparisons were performed with unpaired parametric t test using GraphPad Prism v10.2.3. ANOVA were performed to compare gene expression levels across ethanol concentration treatment groups, with Dunnett’s post-hoc comparisons.

## AUTHOR CONTRIBUTIONS

Conceptualization, C.K., R.C.C., G.T.; methodology, C.K., N.N., E.Y.; formal analysis, C.K., N.N., E.Y.; resources, C.K.; data curation, C.K., N.N., E.Y.; writing—original draft preparation, R.C.C., C.K., N.N., E.Y.; writing—review and editing, R.C.C., C.K., N.N., G.T., E.Y.; funding acquisition, C.K.; All authors have read and agreed to the published version of the manuscript.

## DISCLOSURES

Authors declare no financial or competing interest.

## Notes

### Competing Interest Statement

The authors have declared no competing interest.

